# Evaluating Mycorrhizal association of *Funneliformis mosseae* and *Glomus macrocarpum* with *Solanum melongena L.* (brinjal) using proteomics and system biology approaches

**DOI:** 10.1101/2025.03.17.643673

**Authors:** Subhesh Saurabh Jha

**Affiliations:** Department of Botany, Institute of Sciences, Banaras Hindu University, Varanasi, Uttar Pradesh, Pin-221005

**Keywords:** AMF, *Funneliformis mosseae*, *Glomus macrocarpum*, *Solanum melongena*

## Abstract

Arbuscular mycorrhizal fungi (AMF) symbiotically associate with plant roots and enhance the uptake of inorganic nutrients, increase the ability to withstand the biotic and abiotic stresses. The interaction between plants and microbes can be studied using proteomic. The proteomic profile of an organism can be used to determine the subcellular localization, different chemical environments, potential interaction partners etc. The study was carried out to explore the comparative proteomics of Solanum melongena upon association with *Glomus macrocarpum* and *Funneliformis mosseae*. The study revealed that in comparison to the untreated control, mycorrhizal fungi, *G.macrocarpum* and *F. mosseae* led to upregulation of 43 and 31 proteins respectively. Of these proteins, 26 proteins were common in both the species while 17 were specific to *G. macrocarpum* and 5 were specific to *F. mosseae.* These proteins were predicted to be involved in major physiologically important pathways.

## Introduction

***Solanum melongena*** L., a member of the nightshade family Solanaceae commonly known as brinjal vernacular ‘*baingan’.* Brinjal native to India or South and East Asia and is cultivated worldwide and is a part of multi continent cuisine (Hirakawa et al., 2014). Brinjal is among the top five vegetable crops most valuable for mankind with estimated global production of 54,077,210 metric tonnes annually. Low calorific value and presence of essential micronutrients made brinjal a popular vegetable crop across continents (Watanabe et al., 2016). Brinjal is a rich source of vitamins, especially vitamin K and vitamin B6 (Gürbüz et al., 2018). Among micronutrients brinjal mainly provides potassium, manganese etc. (Watanabe et al., 2016). Secondary metabolites produced by brinjal attributes to its medicinal properties (Vierheilig et al., 1998, Friedman,2015). As per ancient texts it has properties to cure flatulence disorders (Raj et al., 2018). Further metabolites of Brinjal have proven antioxidant and anti-inflammatory properties (Umamageswari and maniyar, 2015).

Arbuscular mycorrhizal fungi (AMF) symbiotically associate with the roots of a number of plants of agricultural interest. More than 85% of all plant families are known to show symbiotic associations with AMF (Jha and Songachan, 2020). This symbiotic relation is beneficial for plants in terms of enhanced uptake of inorganic nutrients especially phosphorus which faces the problem of low-mobility (Ouhaddou et al., 2022). Further AMF are also known to impart ability to withstand various biotic and abiotic stresses viz., resistance to pathogen infections, salinity and drought tolerances etc. This association led to reduction in usage of conventional chemical fertilizers and pesticides which not only improves the economy of agriculture but also protects the environment and food chain from harmful chemicals (Rubin and Gorres, 2020; Silva et al.,2023).

Due to the transgenerational success of mycorrhizal plants that endure repeated harsh conditions, as opposed to that of the less successful non-mycorrhizal counterparts, there is a largely positive belief that mycorrhizas might expedite the evolutionary improvements in plants (Cosme, 2023). Mycorrhizal association with brinjal has a positive impact on crop development and yield, more specially under stress conditions including drought, low nutrient availability, and salinity (Chaturvedi et al.,2018, Malhi et al.,2021).

Mycorrhizal fungi and plants form an advantageous symbiosis that causes a profound restructuring of plant metabolism that involves the control of numerous molecular processes, many of which are still poorly understood (Wang et al., 2023). Proteomics is a potent technique for examining changes associated with interactions between plants and microbes. Further genetic diversity of plants also determines the response of mycorrhizal symbiosis (Zhang et al., 2023).

Proteomics involves the in-depth analysis of the cellular functions, relationships, composition, and architectures of proteins. The proteomic profile of an organism can be used to determine the subcellular localization, different chemical environments, potential interaction partners etc. Eukaryotic cells have a rather complicated proteome that has a wide dynamic range but also has the ability to distinguish between two cellular physiological states (Dupree et al., 2020). Proteomics is practically intricate because it includes the analysis and categorization of overall protein signatures of a genome. Proteome-wide analyses of spatial cellular regulation have been made possible by recent significant advancements in high-throughput microscopy, quantitative mass spectrometry, and interactomics mapping, as well as machine learning tools for data processing (Manes and Nita-Lazar,2018, Sengupta et al.,2021).

Advancement in mass spectrometry coupled Liquid chromatography provides data utilizable in the form of proteogenomics, interactomics, kinomics, and biological pathway modeling through systems biology approach. Comprehensive characterisation of biological pathways for realistic route modelling at the molecular interaction level is a key aim of systems biology (Zhu et al., 2023). These simulations will further help in pathway engineering for crop improvements. Plants and AMF interactions offer a wide variety of benefits but its mechanics are still poorly understood. Here, to understand plant-mycorrhizal association we treated *Solanum melongena* with two mycorrhizal fungi *G. macrocarpum* and *F. mosseae* and studied the effect of their interaction on proteome of *Solanum melongena* plant.

## Material and Methods

### Study site and Biological samples

The study was conducted at botanical garden, department of botany, Banaras Hindu University, Varanasi, India located at 25° 16’ 4.3608’’ N and 82° 59’ 25.7784’’ E with an altitude of 81 meters above the sea level.

### Inoculation of *Solanum melongena* L. plants

For the purpose, pot culture experiment was set up. Control and both the treatments were set up in triplicates. Pure culture of *Glomus macrocarpum* and *Funneliformis mosseae* was obtained by monoculturing both these isolates of AMF. For biofertilizer experiment brinjal (*Solanum melongena* L.) seeds disinfected with 1% sodium hypochloride was placed on sand-soil substrate (1:1/v/v) in 200 ml disposable plastic containers. The germination set up were kept in B.O.D. incubator at 25 under white fluorescent tubes (photoperiod 12 hrs) and watered whenever required to keep the soil mixture moist. After one-month, germinated seedlings of brinjal were transferred in 8X7.6-inch clay pots containing sterilized sand-soil substrate (1:1 v/v) with 10 gms each of both inoculants. Un-inoculated (control) plants were also maintained. AMF colonisation was ascertained by advance microscopy.

### Raising of monosporal cultures of *Glomus macrocarpum* and *Funneliformis mosseae* isolates of AMF

For this study monosporal cultures of both *F. mosseae* and *G. macrocarpum* was used as bio-inoculant. Monosporal culture of both these isolates were obtained in our own lab after investigation and identification of these spores using microscopic techniques. For monosporal cultures of these spores terragreen and sand were used as substrates and *Sorghum bicolor* was used as the host plant. Micropipette tips containing substrate (Terragreen and sand in the ratio of 1:1). 3-4 host seeds (*Sorghum bicolor*) were placed over the substrate. Selected healthy spores of *G. macrocarpum* and *F. mosseae* were kept with the host so as to ensure colonization of germinated spores onto the host. The micropipette tray box containing the culture was filled with water so that the micropipette tips is moistured and incubated in the growth chamber. After the emergence of seedlings from the tips, the tips were taken for drying and this cycle was repeated thrice so as to stimulate vigorous root production. The micropipette tips were then chopped off from below carefully and the seedling were transferred on to big sized pots. The seedling were then allowed to grow and complete its life-cycle (which is about 3-4 months). Following which, the above ground portion of the plant was cut off and the soil and rhizospheric roots were used as pure monosporal culture of *G.macrocarpum* and *F.mosseae* for this study.

### Sample collection and proteome extraction

To purify the proteins from the shoot of control and treated brinjal plants, the TCA-acetone precipitation method was used (Damerval et al.,1986, Carpentier et al., 2005). The obtained protein pellet was cleaned by rinsing twice with ice-cold acetone supplemented with 0.07% β-mercaptoethanol. In between rinsing, samples were incubated for 60 min at −20◦C. Protein pellets were air dried, resuspended in 100 μL sample buffer (50 mM Tris-HCl pH 8.8, 7 M Urea, 2 M Thiourea and 4% CHAPS), and vortexed for 1 hour at room temperature. Final solutions were used for quantification with the BCA method and further processing for LC-MS.

### Sample Preparation

2 ug of protein sample was used for digestion. The sample was first denatured in 8M urea and then reduced with five mM TCEP (Tris[2-carboxyethyl]-phosphine) and further alkylated with 50 mM iodoacetamide (Lopez-Ferrer et al., 2008). The denatured protein samples were subjected to digestion with Trypsin (1:50, Trypsin/lysate ratio) overnight at 37 °C. Digests were dried with a speed vacuum after being cleaned with a C18 silica cartridge to get rid of the salt. The dried pellet was resuspended in buffer A (2% acetonitrile, 0.1% formic acid).

### TMT Analysis

Each (2 μg of) protein sample was supplemented with 2 µl of 1 μg/µl trypsin and 500 µl of 100 mM TEAB buffer and subjected to TMT analysis as described essentially following Wang et al., (2022).

### Database search and data analysis

The mass spectra were hit against the Uniprot protein database as described by Wang et al., (2022) to understand the AMF associations.

## Results and Discussion

### Mycorrhizal Colonization

AMF colonization was observed in all the plantlets inoculated with both the fungi. Plantlets showing the presence of fungal structures were chosen for further proteomic analysis.

### Brinjal leaf Proteome

LCMS profile and peptide fingerprinting analysis showed the presence of 86 different types of proteins (76 high confidence and 10 medium confidence) in leaf proteome of untreated plants. Most proteins reported in control leaf proteomes are involved in normal metabolic processes, protein synthesis and transport pathways. Further treatment of Mycorrhizal fungus *Glomus macrocarpum and Funneliformis mosseae* resulted in production of relatively large numbers of proteins compared to untreated control. *Glomus macrocarpum* treated brinjal leaves proteome contain 113 proteins (104 high confidence & 9 medium confidence) whereas *Funneliformis mosseae* treated brinjal leaves proteome contain 101 proteins (93 high confidence, 7 medium confidence & 1 low confidence).

Proteomics of eggplants upon colonization by AMF have already been reported by several workers (Pasbani et al. 2020). The colonization of AMF with plants is of high significance, since the interaction led to a number of positive changes in the host plant which include ability to tolerate stress (Rodriguez et al. 2008; Ahanger et al. 2014; Salam et al. 2017), and enhanced photosynthetic rate (Birhane et al., 2012). In addition, symbiosis with AMFs improves the plant nutrient uptake and improves in overall plant health (Rouphael et al. 2015; Zou et al. 2016; Thirkell et al. 2017).

Comparative proteomic study revealed that each sample has some unique proteins. Untreated controls have 16 unique proteins which are absent in mycorrhiza treated brinjal leaves. *Glomus macrocarpum* colonization of the *S. melongena* led to induction of 43 proteins in addition to the untreated *S. melongena* proteins, while *Funneliformis mosseae* colonization led to the induction of 31 additional proteins.

**Figure.**
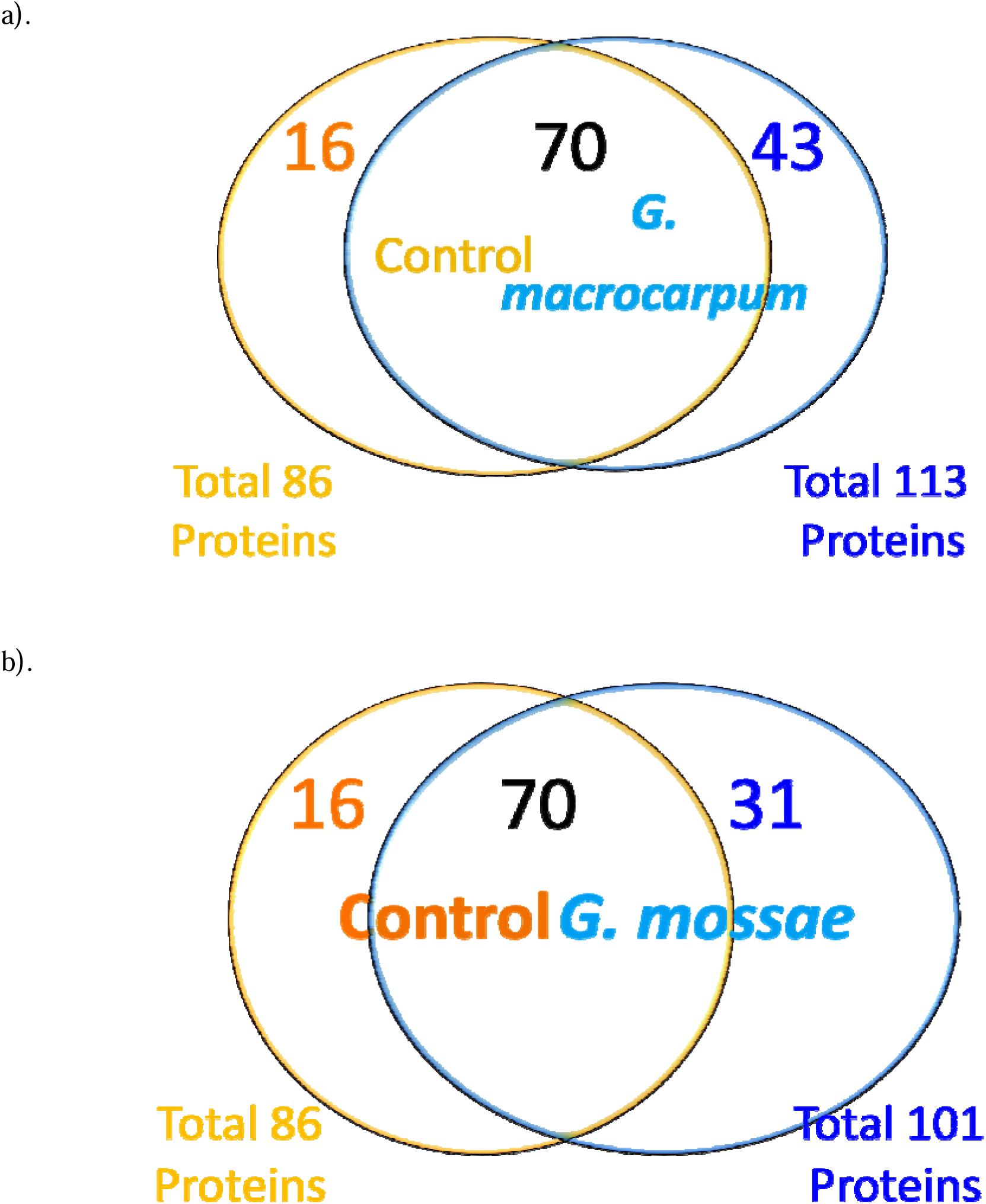
Proteins induced in *Solanum melanogena* upon association with (a) *Glomus macrocarpum*, and (b) *Glomus mosseae*.

**Figure.**
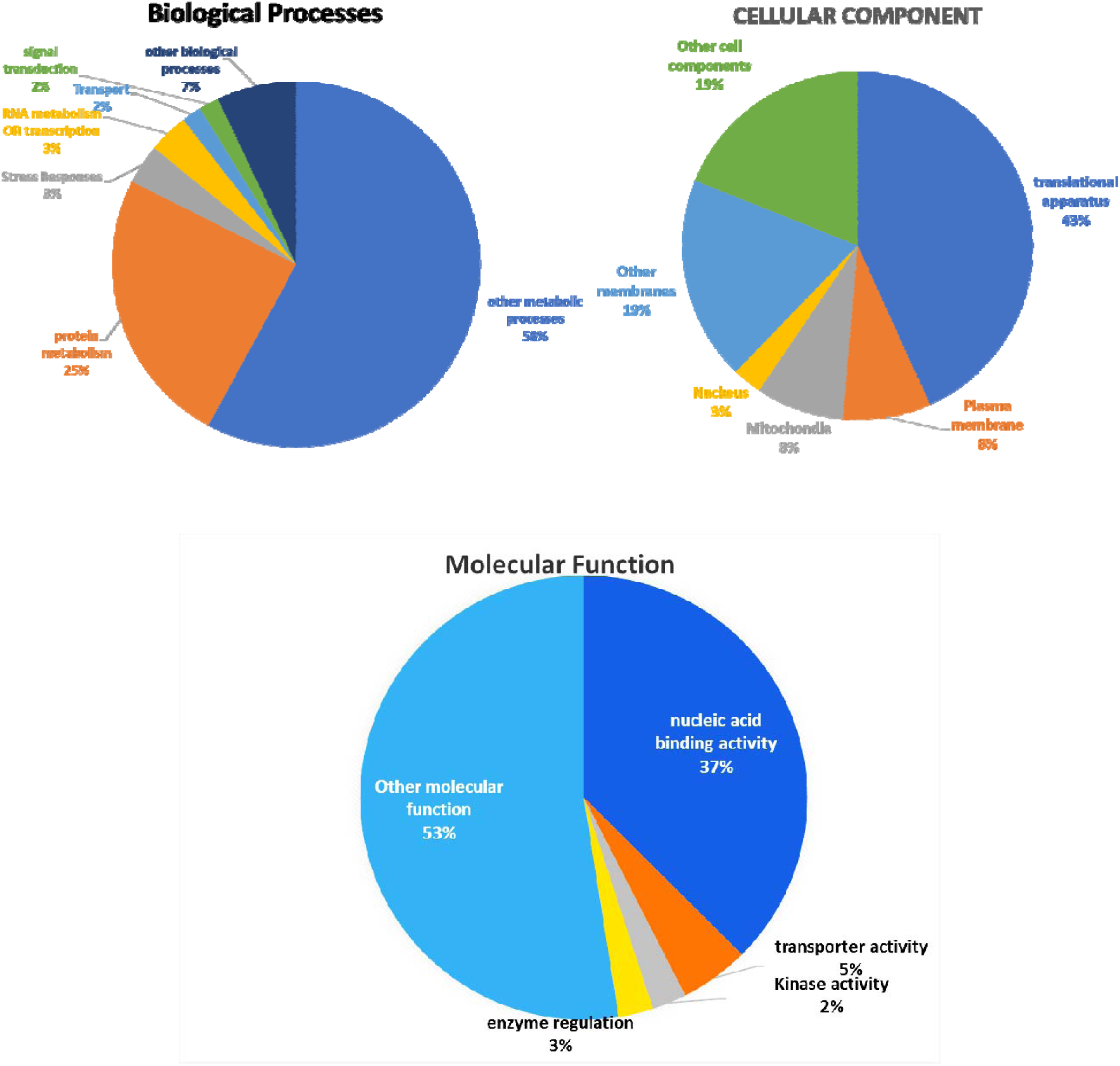
David analysis of the Proteins induced in *Solanum melanogena* upon association with *Glomus macrocarpum*.

### *Glomus macrocarpum* colonization-inducible proteins

David analysis for the biological processes of the proteins induced upon *G. macrocarpum* colonization revealed that one-fourth of the total induced proteins were involved in protein metabolism and, 3% each of the proteins induced were involved in the responding to abiotic stresses and RNA metabolism, while 2% each were involved in signal transduction and transport. 58% of the induced proteins were involved in other metabolic processes and 7% in other biological processes. Majority of the proteins were located in translation apparatus (43%), followed by plasma membrane (8%), mitochondria (8%) and nucleus (3%). While 19% of the proteins were located in other membranes and other 19% in other cell components. Nearly 37% of the *G. macrocarpum* inducible proteins were involved in the nucleic acid binding activity, while a small fraction i.e., 5%, 3% and 2% of the proteins were involved in the transporter activity, enzyme regulation and kinase activity. Most of the protein i.e., 53% were involved in other molecular functions.

### *Glomus mosseae* colonization-inducible proteins

David analysis for the biological processes of the proteins induced upon *G. mosseae* colonization revealed that 32%, 8%, and 2% of the *G. mosseae*-inducible proteins were involved in Protein metabolism, transport and stress response. 54% of the proteins were involved in other metabolic processes while 4% were involved in other biological processes. Forty-six percent proteins were part of translational apparatus, and 9% and 3% were located in plasma membrane and mitochondria respectively. Whereas, 18% of the proteins were located in other membranes and 24% were present in other cell components. 34% of the *G. mosseae* colonization induced proteins were implicated in the nucleic acid machinery and 6% and 2% were implicated in the transporter and kinase activity.

The association of the AMF, *G. macrocarpum* with the eggplant led to induction of more number of proteins than that of the *F. mosseae*. RNAseq analysis of the eggplant in association with the two AMFs, i.e., *G. macrocarpum* and *F. mosseae* also revealed high transcriptional activity in former than the latter (Subhesh Saurabh Jha and L.S.Songachan, communicated elsewhere). Detailed analysis of these proteins for biological processes, cellular component and molecular functions revealed that the induction of proteins involved in the biological processes, signal transduction and RNA metabolism, those belong to cellular component, nucleus, and molecular function of enzyme regulation were found only in the species *G. macrocarpum*. Further, efforts were also made to analyse the AMF inducible proteins of *S. melongena* in response to both *G.macrocarpum* and *G. mosseae*. The proteins expressed in response to different AMFs may vary from species-to-species. These changes might be contributing to the factors regulating the progression of symbiotic association between the fungus and host plant (Begum et al., 2019). Further efforts were also made to closely look at the AMF inducible proteins.

**Figure.**
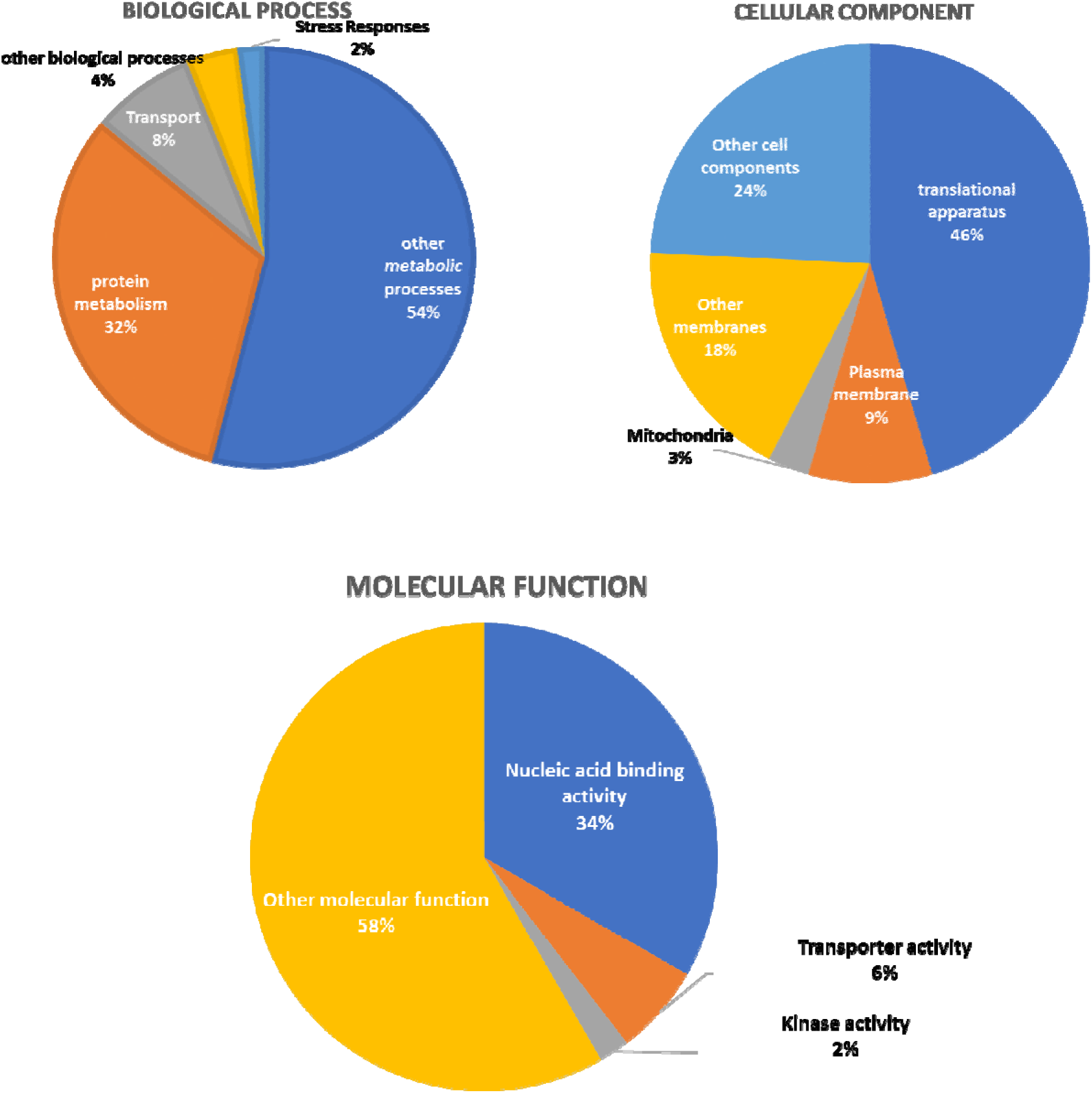
David analysis of the proteins induced in *Solanum melanogena* upon association with *Glomus mosseae*.

### Comparison of Proteins induced in response to *Glomus macrocarpum* or *G. mosseae*

Association of both the AMFs with *S. melongena* resulted in induction of 26 proteins, while 17 proteins were specific to *G. macrocarpum* and 5 proteins were specific to the association with *G. mosseae* with *S. melongena* (Figure ).

**Figure:**
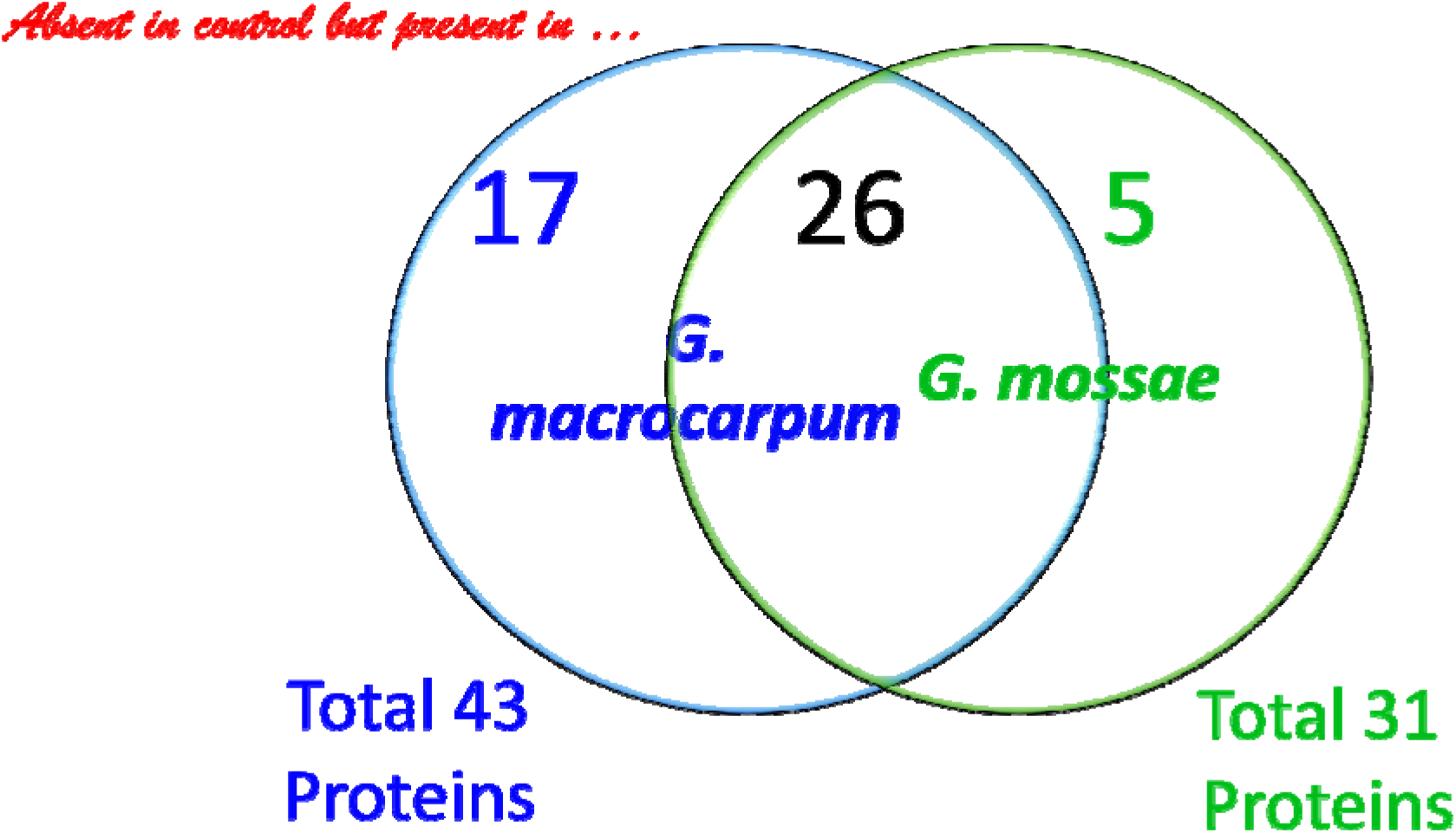
Proteins specific to the colonization of *Glomus mosseae*, and *Glomus macrocarpum*.

### Mycorrhizal Colonization associated proteins

Association of the *Solanum melongena* with both the *Glomus* sp. induced 26 proteins (23 high confidence and 3 medium confidence). These induced proteins are involved in protein synthesis (Ribosomal protein Subunit S2, S3, S4, S7, S8, S12, S14, S16, S18, S19, L16, L20, L22, L33), biosynthesis of the polyphenol and flavonoids (Phenylalanine ammonia-lyase, Flavanone 3-hydroxylase), respiratory (NADH-quinone oxidoreductase subunit D, Subunit I, NADH dehydrogenase [ubiquinone] iron-sulfur protein 3), signaling (Inositol-1-monophosphatase), Transferase (Aminotransferase, Acetyl-coenzyme A carboxylase carboxyl transferase subunit beta), photosystem proteins (Photosystem II reaction center protein H), stress responders (I2 (Fragment), and Protein kinase domain-containing protein (Table 1).

**Table 1:**
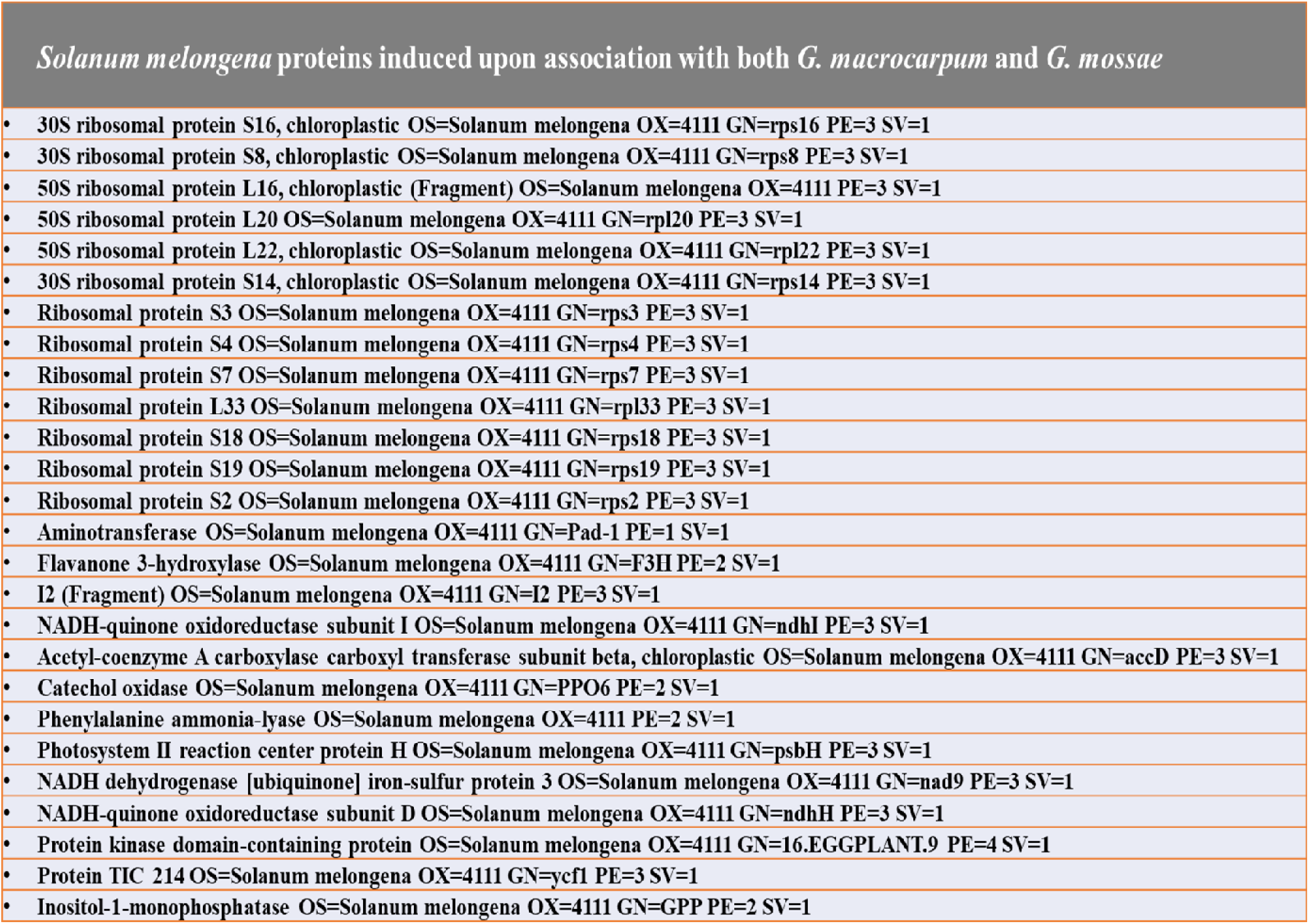
Proteins expressed in the *Solanum melongena* in association with both the AMFs, *G. macrocarpum* and *G mosseae*.

### *Glomus macrocarpum*-specific induced proteins of *S. melongena*

A total of seventeen proteins were induced upon association of the *S. melongena* with *G. macrocarpum* (Table 2). The seventeen proteins included proteins involved in flavanoid biosynthesis (Chalcone-flavonone isomerase, Flavonol synthase), Cytochromes (Cytochrome b6, Cryptochrome 1, Cytochrome P450 77A1 and DNA photolyase protein), transcription related proteins (DNA-directed RNA polymerase, NAC1 transcription factor, Transcription repressor, Maturase-related protein and 4-coumarate:coenzyme A ligase), stress responders (BURP domain-containing protein, DUF4338 domain-containing protein), glycoproteins (Miraculin homologue, Anthocyanidin 3-O-glucosyltransferase), Ascorbate synthesizing enzyme (L-galactono-1,4-lactone dehydrogenase) and a starch synthase enzyme (Granule-bound starch synthase) (Table).

**Table 2:**
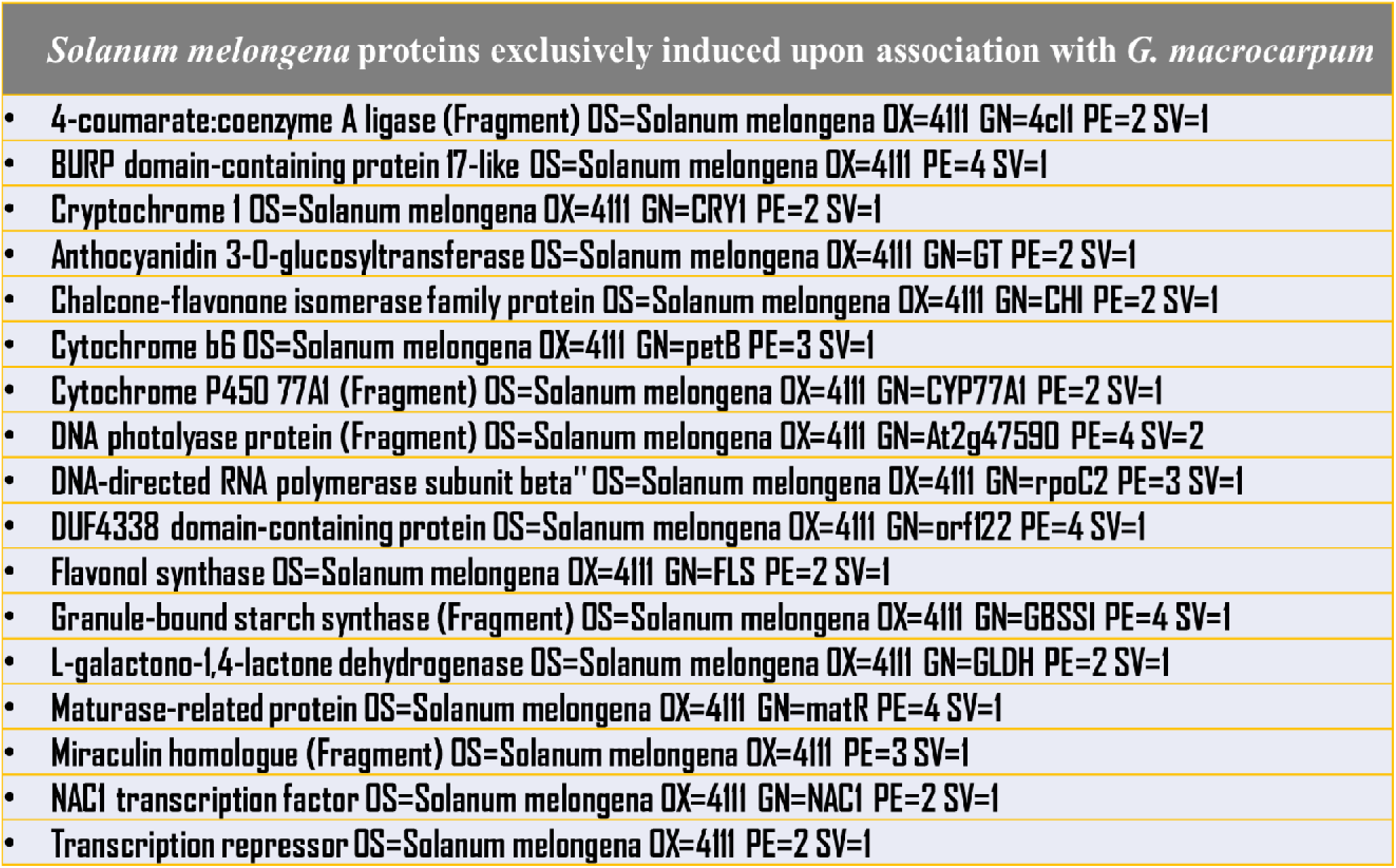
Proteins exclusively expressed in the *Solanum melongena* in association with *G. macrocarpum*.

| *Solanum melongena* proteins exclusively induced upon association with *G. macrocarpum* |
| --- |
| • 4-coumarate:coenzyme A ligase (Fragment) OS=Solanum melongena OX=4111 GN=4c11 PE=2 SV=1 |
| • BURP domain-containing protein 17-like OS=Solanum melongena OX=4111 PE=4 SV=1 |
| • Cryptochrome 1 OS=Solanum melongena OX=4111 GN=CRY1 PE=2 SV=1 |
| • Anthocyanidin 3-O-glucosyltransferase OS=Solanum melongena OX=4111 GN=GT PE=2 SV=1 |
| • Chalcone-flavonone isomerase family protein OS=Solanum melongena OX=4111 GN=CHI PE=2 SV=1 |
| • Cytochrome b6 OS=Solanum melongena OX=4111 GN=petB PE=3 SV=1 |
| • Cytochrome P450 77A1 (Fragment) OS=Solanum melongena OX=4111 GN=CYP77A1 PE=2 SV=1 |
| • DNA photolyase protein (Fragment) OS=Solanum melongena OX=4111 GN=At2g47590 PE=4 SV=2 |
| • DNA-directed RNA polymerase subunit beta'' OS=Solanum melongena OX=4111 GN=rpoC2 PE=3 SV=1 |
| • DUF4338 domain-containing protein OS=Solanum melongena OX=4111 GN=orf122 PE=4 SV=1 |
| • Flavonol synthase OS=Solanum melongena OX=4111 GN=FLS PE=2 SV=1 |
| • Granule-bound starch synthase (Fragment) OS=Solanum melongena OX=4111 GN=GBSSI PE=4 SV=1 |
| • L-galactono-1,4-lactone dehydrogenase OS=Solanum melongena OX=4111 GN=GLDH PE=2 SV=1 |
| • Maturase-related protein OS=Solanum melongena OX=4111 GN=matR PE=4 SV=1 |
| • Miraculin homologue (Fragment) OS=Solanum melongena OX=4111 PE=3 SV=1 |
| • NAC1 transcription factor OS=Solanum melongena OX=4111 GN=NAC1 PE=2 SV=1 |
| • Transcription repressor OS=Solanum melongena OX=4111 PE=2 SV=1 |

David analysis of the proteins commonly induced in *S. melongena* upon association with *G. macrocarpum* and *G. mosseae*. Out of these proteins, 33% were involved in protein metabolism, 5% in transport, and 2% in stress response. 55% of the proteins were involved in other metabolic processes and 5% in other biological processes. Analysis of cellular component revealed that 36% of the proteins belonged to translational apparatus, 4% and 11% in plasma and other membranes respectively, and 2% belonged to mitochondria, while 47% belonged to other cell components. 7% of these proteins were involved in transporter activity and 4% were involved in kinase activity, whereas, rest 89% perform other molecular functions.

**Figure.**
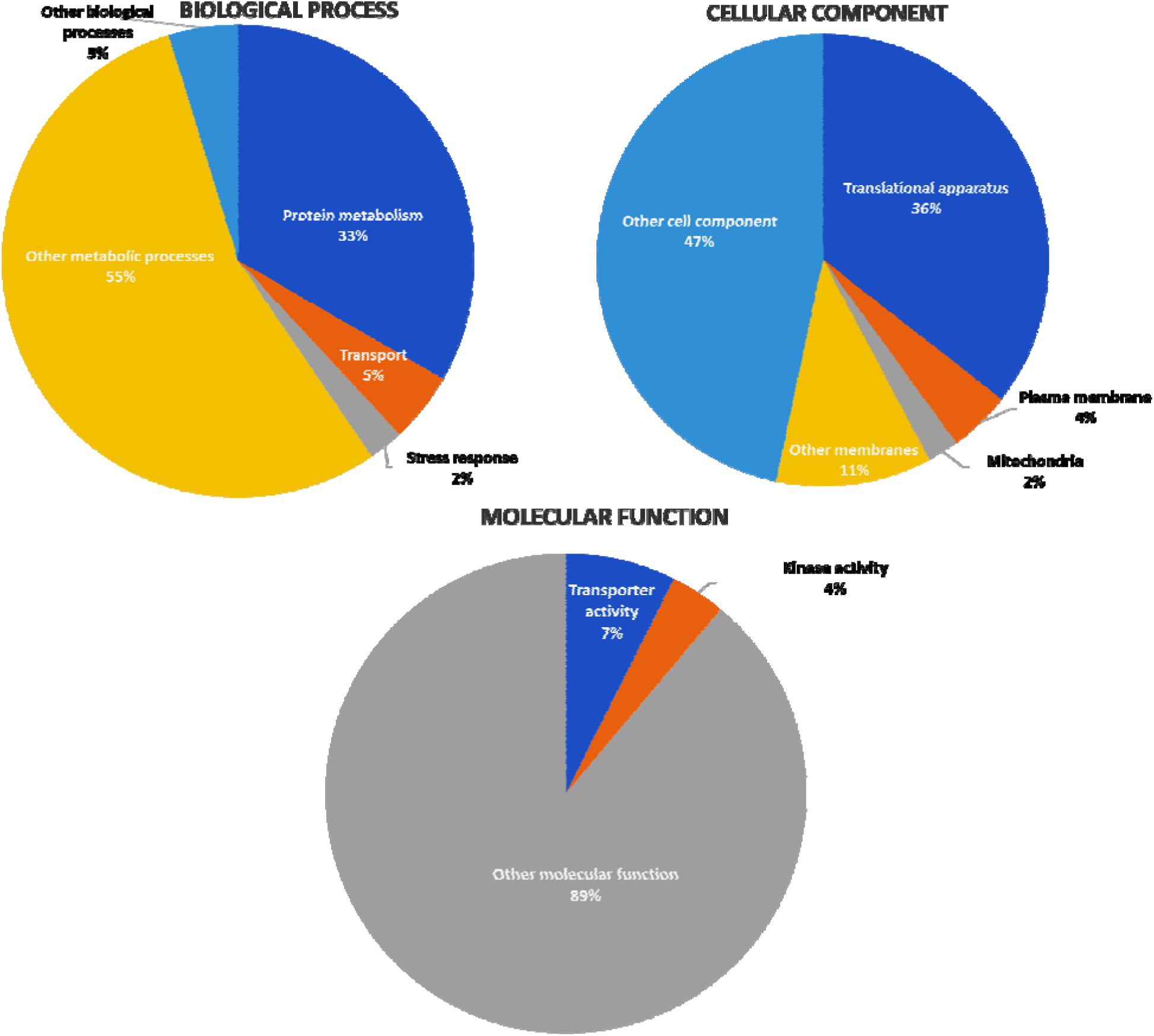
David analysis of the Proteins induced in *Solanum melanogena* upon association with *Glomus macrocarpum* and *Glomus mosseae*.

The proteins specific for *S. melongena - G. macrocarpum* association comprised of proteins involved in RNA metabolism (12%), Stress response (6%) and Signal transduction (6%). 59% of the proteins were involved in other metabolic processes and 17% were involved in other biological processes. 22% of the proteins were belonged to mitochondria, 11% to nucleus, 45% to membranes other than the plasma membrane and 22% to other cell components. 15% of the protiens had nucleic acid binding activity, 8% had enzyme regulator activity while 77% had other molecular functions.

### *Glomus macrocarpum*-specific induced proteins of *S. melongena*

Five proteins were found to be inducible in the *S. melongena* exclusively upon association with *G. mosseae* (Table 3). The five unique proteins are known to be involved in flavanoid biosynthesis (Catechol oxidase, NADH-quinone oxidoreductase subunit B), translation (30S ribosomal protein S11, 50S ribosomal protein L32) and stress response (MYBL1).

**Table 3:** Proteins exclusively expressed in the *Solanum melongena* in association with *G. mosseae*.

| *Solanum melongena* proteins exclusively induced upon association with *G. mosseae* |
| --- |
| • 30S ribosomal protein S11, chloroplastic OS=*Solanum melongena* OX=4111 GN=rps11 PE=3 SV=1 |
| • 50S ribosomal protein L32, chloroplastic OS=*Solanum melongena* OX=4111 GN=rpl32 PE=3 SV=1 |
| • Catechol oxidase OS=*Solanum melongena* OX=4111 PE=3 SV=1 |
| • MYBL1 OS=*Solanum melongena* OX=4111 GN=MYBL1 PE=4 SV=1 |
| • NADH-quinone oxidoreductase subunit B OS=*Solanum melongena* OX=4111 GN=ndhK PE=3 SV=1 |

David analysis for biological processes revealed that one-fourth of the proteins were involved in protein metabolism, and 12% were involved in transport, whereas, 63% were involved in other metabolic processes. One-fourth of the proteins were located in translational apparatus. Another one fourth were present in plasma membrane (12%) or other membranes (8%). Rest half of the proteins were present in other cell components. The molecular function analysis revealed that 29% of the proteins were involved in nucliec acid binding, 14% in the transporter activity while rest 57% were involved in other molecular functions.

**Figure:-.**
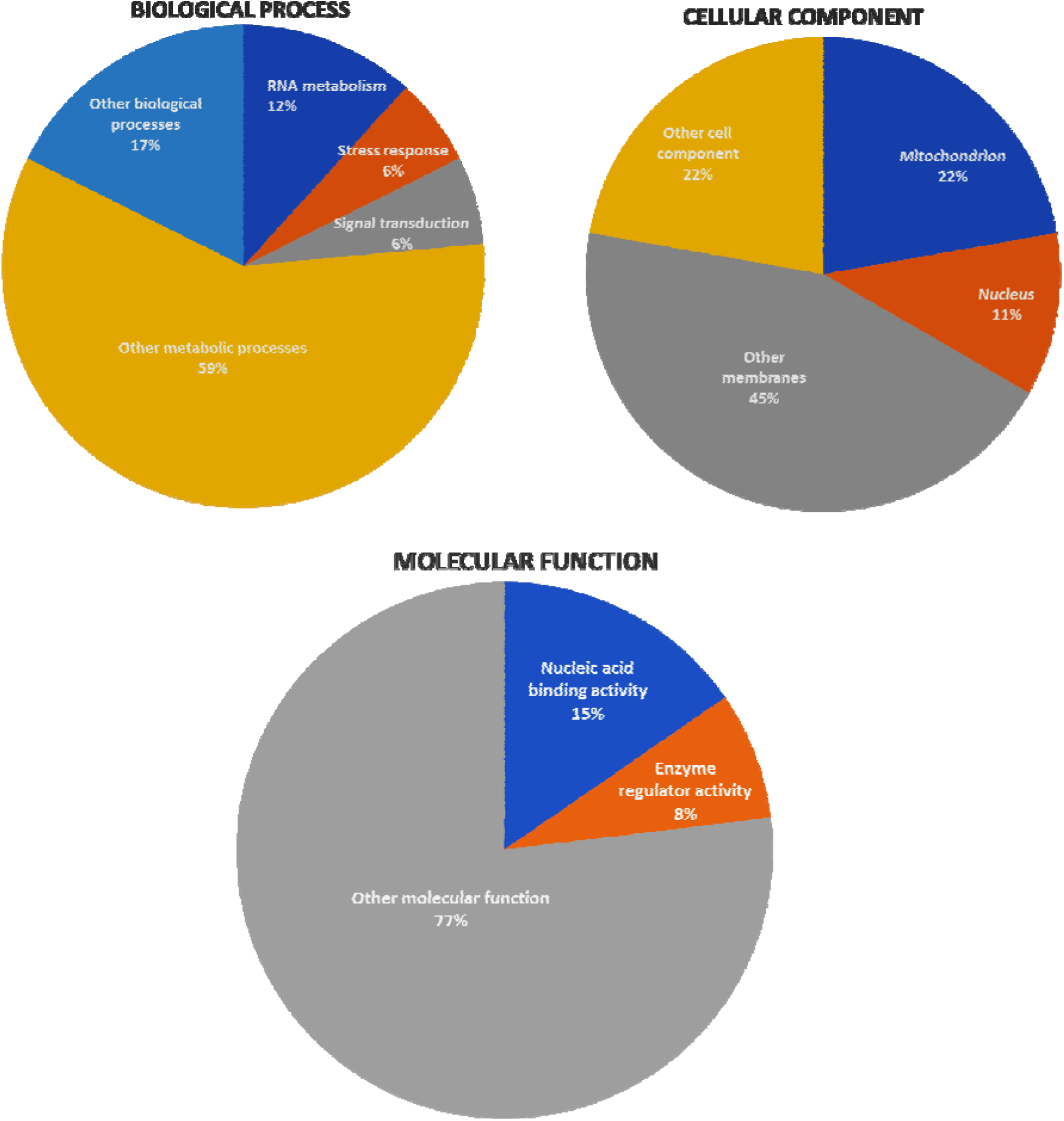
David analysis of the Proteins induced exclusively in *Solanum melongena* upon association with *Glomus macrocarpum*.

The comparisons of the *G. macrocarpum* specific and *G. mosseae* specific inducible proteins of S. melongena revealed induction of the 6 proteins involved in the signal transduction and 12 involved in RNA metabolism only in the case of association with *G. macrocarpum*. Moreover,, 6 additional proteins involved in the stress management were observed in *S.melongena*- *G.macrocarpum* association. It is worth mentioning overhere that high induction of anti-stress genes at transcript levels were also observed in the case of association with *G. macrocarpum* with *S. melongena* but not in the case of *G. mosseae* association. The transcripts that were seen upregulated in the study were also found to express at protein levels. However, the physiological relevance of the study need to be validated in future studies.

In conclusion, *G. macrocarpum* and *G. mosseae* are two important arbuscular mycorrhizal fungi that form symbiotic relationship with the eggplant, *Solanum melongena*. The symbiotic association with the fungi trigger proteomic changes in the plant. A lot of proteins are induced in response to the association with both the fungi, yet the response to *G. macrocarpum* is more pronounced than that of the *G. mosseae*. The AMF-specific protein factors are candidates for future research in this field and may increase out understanding fo the symbiotic association between the plant and the fungi (Begum et al., 2019). The physiological aspect of these interactions and functional relevance of these proteomic changes are also subject of further research.

**Figure.**
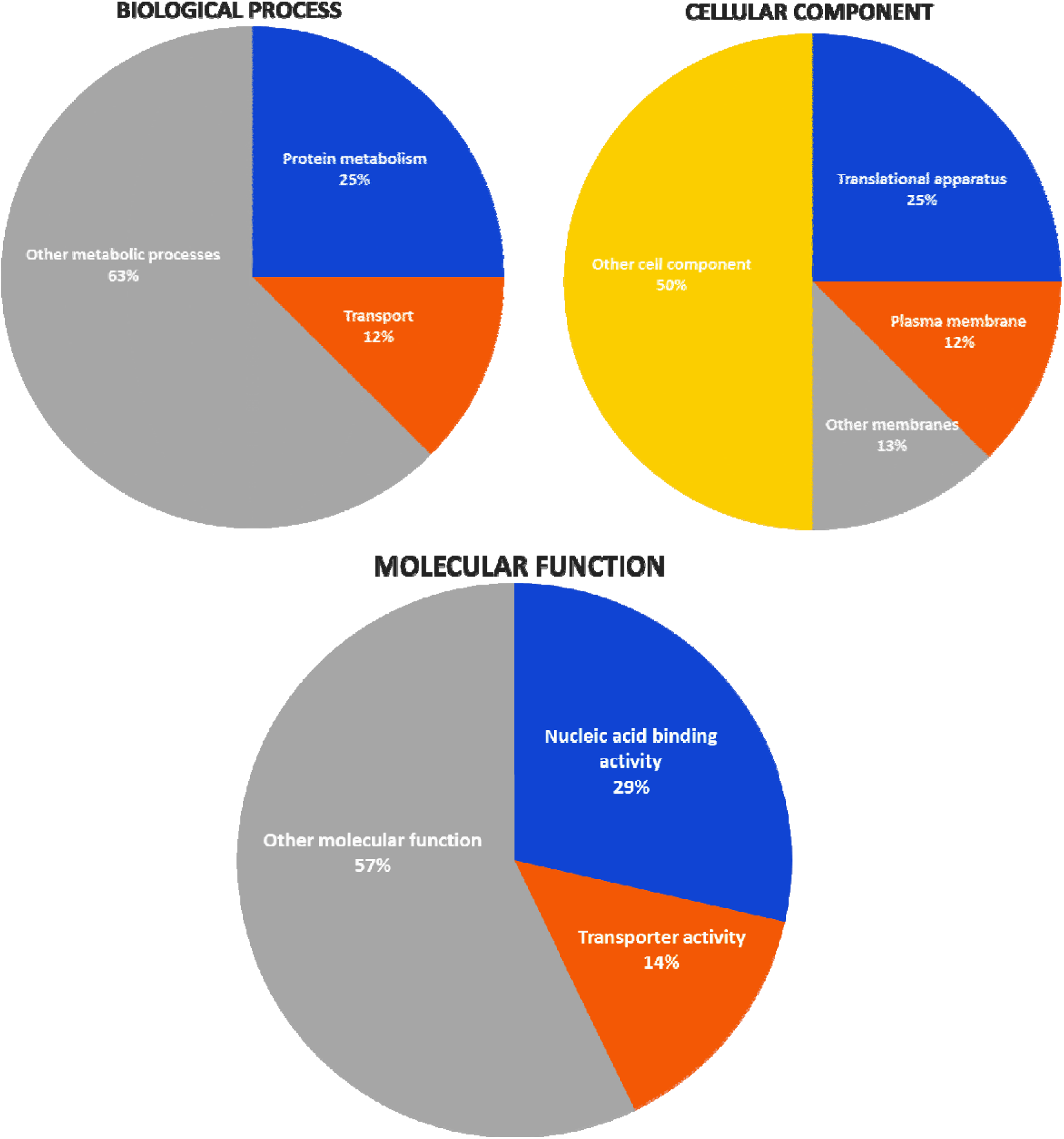
David analysis of the Proteins induced exclusively in *Solanum melanogena* upon association with *Glomus mosseae*.

